# CATHe2: Enhanced CATH Superfamily Detection Using ProstT5 and Structural Alphabets

**DOI:** 10.1101/2025.06.22.660903

**Authors:** Orfeú Mouret, Jad Abbass

## Abstract

**Motivation:** The CATH database is a free publicly available online resource that provides annotations about the evolutionary and structural relationships of protein domains. Due to the flux of protein structures coming mainly from the recent breakthrough of AlphaFold and therefore the non-feasibility of manual intervention, the CATH team recently developed an automatic CATH superfamily classifier called CATHe, that uses a feed-forward network classifier with protein Language Model (pLM) embeddings as input. Using the same dataset, in this paper, we present, CATHe2 that improves on CATHe by switching the old pLM ProtT5 for one of the most recent versions called ProstT5, and by introducing domain 3D information as input to the classifier, in the form of Structural Alphabet representation, namely 3Di sequence embeddings. Finally, CATHe2 implements a new version of the feed-forward network (FNN, i.e, non-recurrent neural network) classifier architecture, fine-tuned to perform at the CATH superfamily prediction task.

**Results:** The best CATHe2 model reaches an accuracy of 92.2 ± 0.7% with an F1 score of 82.3 ± 1.3% which constitutes an improvement of 9.9% on the F1 score and 6.6% on the accuracy, from the previous CATHe version (85.6 ± 0.4% accuracy and 72.4 ± 0.7% F1 score) on its largest dataset (~ 1700 superfamilies). This model uses ProstT5 AA sequence and 3Di sequence embeddings as input to the classifier, but a simplified version requiring only AA sequences, already improves CATHe’s F1 score by 6.7 ± 1.3% and accuracy by 6.6 ± 0.7% on its largest dataset.

**Availability & Implementation:** The code is available on https://GitHub.com/Mouret-Orfeu/CATHe2. Datasets: https://doi.org/10.5281/zenodo.14534966

## 1. Introduction

Proteins are essential to life, forming the foundation of virtually all biological processes. They are complex nanomachines with specific functions which largely depend on their unique 3D structure. Figuring out what shapes proteins fold into is known as the “protein folding problem”, and has stood as a grand challenge in biology since 1972 [1, 2]. Alphafold2 is a 2020 automated algorithm which reached experimental performance in the protein folding problem, which is now considered solved for a large part of known proteins [3]. Since then, a database of Alphafold inferred 3D protein conformations also called CSM (Computed Structure Model) has been filled with over 200 million protein structure predictions. This database is the Alphafold database or AFDB for short [4]. As protein evolution gives rise to families of structurally related proteins, within which sequence identities can be extremely low, structure-based classifications are needed to identify unanticipated relationships in known structures, now present in overwhelming numbers. CATH is a classification system that fills this need, and even allows inference of domain function when paired with FunFam subclassification [5]. The corresponding CATH database is updated daily with the release of a snapshot version of the most up-to-date classifications. A full-release version called CATH-Plus including a greater depth of information about CATH domains is also uploaded annually, benefiting from ever improving automatic classification algorithms [6, 7, 8]. The last CATH-Plus version available at the time of writing is 4.4.0 according to the CATH website [9], this version includes 6,573 superfamilies.

After the major bioinformatics revolution with AlphaFold2 (AF2) [3], bioinformaticians are now witnessing a second revolution with the rise of protein Language Models (pLMs) [10, 11, 12, 13]. After the surprising huge leap of performance in NLP (Natural Language Processing) allowed by LLMs (Large Language Models) in recent years, protein processing soon took advantage of this new technique to process Amino Acid (AA) sequences, resulting in automatic and powerful feature extractors such as the ProtT5 pLM and CATHe classifier [14] (in this paper, ‘ProtT5’ refers to ProtT5-XL-U50 from the T5 models presented in the ProtTrans paper [15]). With new pLMs, pLM usage, and larger protein 3D conformation databases, the remote homologue detection and homology based inference (HBI) tool landscape has been rapidly evolving, making use of fast and sensitive large scale remote homologue detection using structural information [16, 17, 18] or better feature extraction from primary sequences.

While standard sequence profile and Hidden Markov Model (HMM)-based alignment tools such as MMSeqs2 [19] and HHsuite [20, 21] have been successful, they reach their limits in the twilight or midnight zones of sequence similarity. In contrast, structural alignment tools like Foldseek [18] leverage 3D protein information to perform better at detecting remote homology, and are significantly faster than earlier 3D alignment methods such as DALI [22]. These techniques are the state of the art in their own category, but the new wave of techniques inspired by pLMs is already being regarded as more promising as they can be faster and more sensitive in some cases. Embedding-based annotation transfer (EAT) transfers protein annotation to a close protein using distance test, like Euclidean distance or cosine similarity between pLM embeddings [23, 24, 25, 26]. EAT can be coupled with Contrastive Learning (CL) to extend the distance between embedding distribution clusters and thus improving EAT results [25, 27]. Protein Embeddings Based Alignment (EBA or PEbA) uses pLM embeddings alignment to find homologues and has been very successful, with a lot of work using it to beat former state-of-the-art methods in various domains [28, 29, 30]. Protein artificial neural network (ANN) classifiers, such as those used in CATHe and this study, are also valuable tools for homology-based inference (HBI). These classifiers perform HBI by using supervised learning to develop internal representations for homology detection.

All of the previous techniques are based on 3 possible types of input. First primary sequences, that can be vectorized in many different ways, pLM embeddings is one of them. Secondly evolutionary information, including Multiple Sequence Alignment (MSA), (HMM based techniques use MSA but some pLM based techniques too [31, 32, 33, 34]), and the last input type is structural information, which can take multiple forms like full and precise 3D shape or just contact information, secondary structure annotation, or 1D representation like with Structural Alphabets (SA) [35]. All this structural information can be inferred information or ground truth and used as is or embedded using pLM for more complex feature extraction [36, 37, 38, 39, 26]. Recent structural alignment tools are promising, potentially more meaningful and accurate for HBI since structure is more important for protein function than primary sequences, plus, most pairs of proteins with similar structures populate the midnight zone [40]. Nowadays, many new techniques revolve around finding shortcuts to use 3D information without having to compute the 3D shape of proteins or skipping MSA construction, and thus speeding up potential subsequent structural alignment [41, 42, 43]. Using both structural and primary sequence information, i.e hybrid-information based methods have been shown to outperform other categories of methods [44] for remote homology detection and HBI. As primary sequences have been already studied extensively current research is focused on making the most out of protein 3D structure. Most sensitive and fast remote homology detection approaches today use embedding-based and structure-aware tools such as Foldseek, ESMFold [45] or AlphaFoldMultimer [46] for instance, to produce relevant data for subsequent HBI techniques like EBA which seems to be one of the most effective. Some techniques like Foldseek-TM [18] use shortcuts to be even faster but less sensitive [28]. However, there is a way to be even faster, without losing in sensitivity, i.e, using discriminative methods [10]. Discriminative methods realize HBI tasks without the need for any kind of alignment, as with the ANN classifiers of CATHe and CATHe2. This is why automated protein annotation with ANN classifiers is a good long-term solution for large scale specific HBI tasks. Although it is less transparent, ANN classifiers are potentially faster and more accurate as no alignment is required and the classifier learns on its own how to find homologs, whatever the complexity of the homology relation. Given the right amount of representative data, sufficiently complex ANN classifiers can learn the proper way to classify (and annotate) proteins based on their most relevant properties extracted by a trained pLM, benefiting from transfer learning.

In this paper, we introduce CATHe2, where both AA sequence embeddings and 3D structure information are used to perform the HBI of CATH annotations with an ANN classifier. The 3D information which is also embedded thanks to the 3Di SA developed for Foldseek and the ProstT5 pLM. The first version of CATHe relied on both pLM AA sequence embedding and ANN classifier usage to improve on the automatic CATH annotation task up to an F1 score of 72.4% on the corresponding dataset. CATHe2 improves on this method, using newer pLMs, adding protein 3D information as input (in the form of SA embeddings) and fine tuning the previous ANN classifier architecture for a final model F1 score of 82.3% on almost the same dataset. But CATHe and CATHe2 are not the only CATH annotation algorithms tested in recent years, many different pLMs were tested with different metrics, across many papers using different techniques and approaches for CATH annotation inference. As shown at the end of the Results paragraph, the current ranking of HBI techniques for CATH annotations evidenced by literature review seems to be classifiers >EBA >EAT+CL >EAT. Similarly, a ranking of pLM tested for this task should resemble ProstT5 >ProtT5 >TM-Vec >Ankh large >ESM2 >ESM1b although this ranking depends strongly on HBI technique. Almost all of these pLMs were also tested for CATHe2.

## 2. Materials and methods

### 2.1. Data and datasets

To conduct as accurate and fair comparison as possible, CATHe2 datasets are based on the CATHe ones [14]. In CATHe, two SF datasets were used. The large SF dataset contains the 1,773 SF which had at least 2 non-redundant corresponding sequences in PDB at the time of CATHe development. This SF dataset represented 87.6% of the CATH v4.3 database in terms of domain number. The smaller SF dataset is a subset of the bigger one, only keeping the most populated 50 SF, representing 37.32% of the CATH v4.3 domain sequences in total. For each SF in the corresponding SF dataset, at least one domain sequence was placed in the test and validation set. Then, training sets were built using CATH-Gene3D sequences corresponding to the respective test and validation set SFs. In addition, sequences from PDB corresponding to the 3,456 SF without 2 non-redundant sequences in PDB were included via the ‘other’ part of the training dataset. All these other SF were under the single SF label ‘other’ in the test and validation set, as there were not enough sequences in PDB to populate the test and validation set with each of these SF.

The sample sizes of the corresponding training datasets are 1,039,135 domains and 528,863 domains for the large and small training set respectively. For reference, CATH database contains 6,573 SF at the time of writing, but most of them are admittedly very rare as, in 2023 the 200 most highly populated superfamilies in CATH alone represent 62% of AF2 domains [7]. The CATHa dataset building process include a sequence identity filter in order to assesse the CATHe model using a dataset of remote homologues having less than 20% sequence identity to any domain in the training set.

Although CATHe datasets were left as is for CATHe2 as much as possible, some of the dataset protein domains were removed to test and train CATHe2 due to the use of 3Di sequences, however, as discussed below, the missing proteins can be seen as negligible when performance is compared. For the small dataset, there was no loss whatsoever. In the large dataset, the inability to find the PDB files to compute 3Di properly for some domains resulted in the loss of 137,698 domains in this dataset, and 32 superfamilies among the 1773 formerly present in this dataset. These lost SF were among the least represented in the training set (please see Figure 1), which implies they were rare SF in the first place. Moreover, the ‘other’ SF representing all other SF than the 1772 present in the CATHe main dataset was not included in the CATHe2 dataset, due to a lack of information on these domains. It is worth mentioning that in the experiment where only AA sequence was considered, the dataset of CATHe2 does not suffer any superfamily loss compared to CATHe.

**Figure 1:**
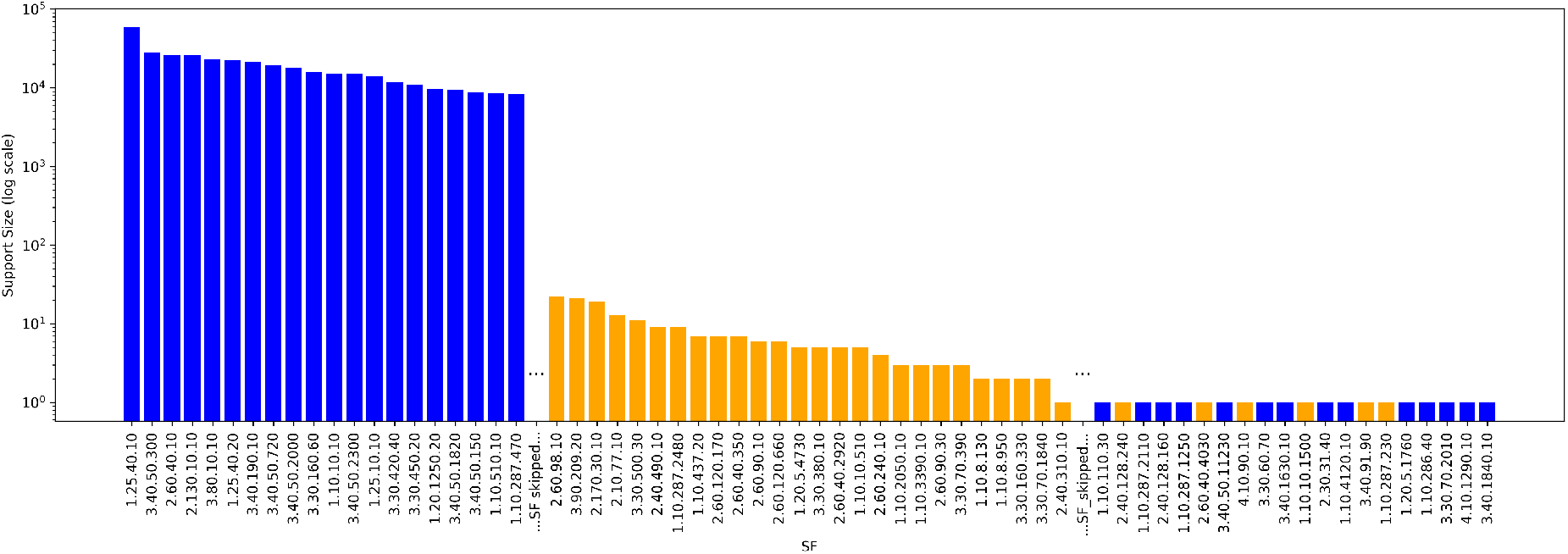
Lost SF by support in CATHe training set. This graph presents the support size of some superfamilies in the CATHe training set, on a logarithmic scale. Orange bars represent lost SF for the 3Di sequence dataset, meaning the SF lost due to unavailability of 3D structure for 3Di computation. As mentioned in Data and datasets, the lost SFs had low support in this dataset.

For the sake of further and advanced experiments, we have created some smaller instances of the CATHe datasets by applying two different filters resulting in CATH SF loss, although none of these filters affected the number of SF lost in the final CATHe2 models. The first filter tested was the predicted local distance difference test (pLDDT) threshold filter. One of the most interesting features generated by AF2 is a confidence score, i.e., pLDDT [47]. Since the accuracy of the CSM directly affects the accuracy of the 3Di sequences and consequently the accuracy of the models, we have tried to remove as much as possible the ‘low quality’ structures in the hope to improve the models. Since the 3Di sequences from CATHe training dataset are computed on CSM from AFDB, which provides a confidence score, namely the pLDDT score, we have computed the overall pLDDT of all domains to be considered in the filter. The pLDDT threshold filter removes domains from the training set for which the pLDDT is under a certain threshold (pLDDT thresholds tested for CATHe2 models are: 0, 4, 14, 24, 34, 44, 54, 64, 74, 84); the SF completely removed from the training set by this filter were also removed from the test and validation set, see Figure 2. Needless to say, a high threshold for such a filter not only leads to loss of SFs but also the reduction of the size of each class representing SFs. The second filter tested was a support threshold. As seen in Figure 1, the size of classes (size called support of the classes, a class being an SF here), is not only quite unbalanced, but also some SFs comprises extremely low number of domains (< 10). This affects the ability of the models to train on such SFs, therefore, a support threshold filter was tested to try removing low support SF and improve the classifier performance. Support thresholds tested for CATHe2 models were 0 and 10, where 0 refers to not applying the filter. The SF completely removed from the training set by this filter were also removed from the test and validation set. A rather high number of SF was lost with support threshold = 10 (these lost SF include 167 SF which had already a support inferior or equal to 10 in CATHe raw training set, and 5 SF for which support was lowered under 10 by unfound 3D structure for 3Di usage). For this reason, no models trained with support threshold 10 have made it to the final version of CATHe2 although performance metrics were higher, see Section 3 for more details. Figure 2 summarises these two filtering processes and Figure 3 shows training set size in domains and SF relatively to the different filter thresholds.

**Figure 2:**
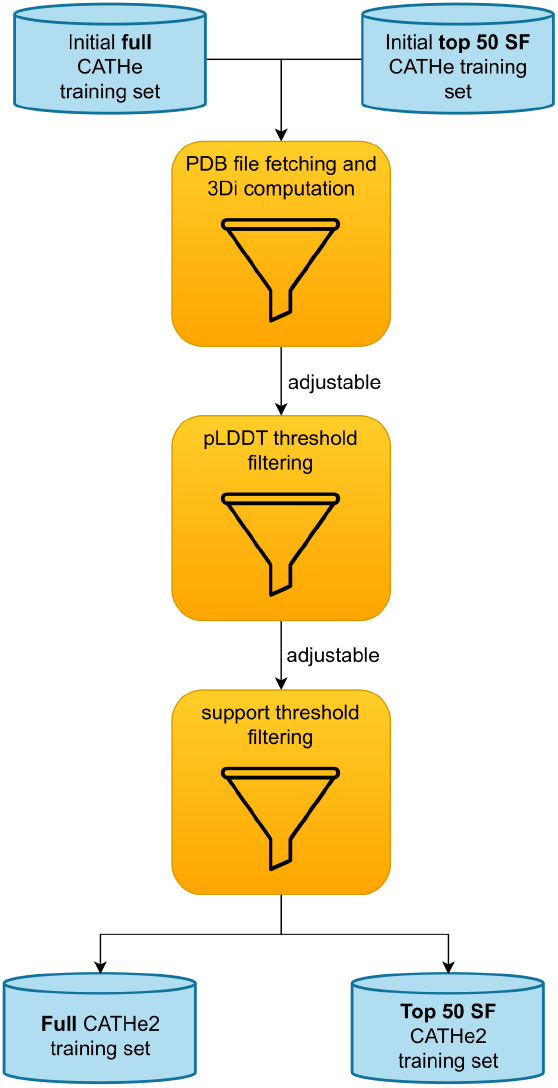
Overview of the dataset filtering process. Filtering process of CATHe2 datasets. Each filter is optional, pLDDT threshold and support threshold have adjustable values and many combinations were tested to improve model performance as much as possible without losing too much SF in the process.

**Figure 3:**
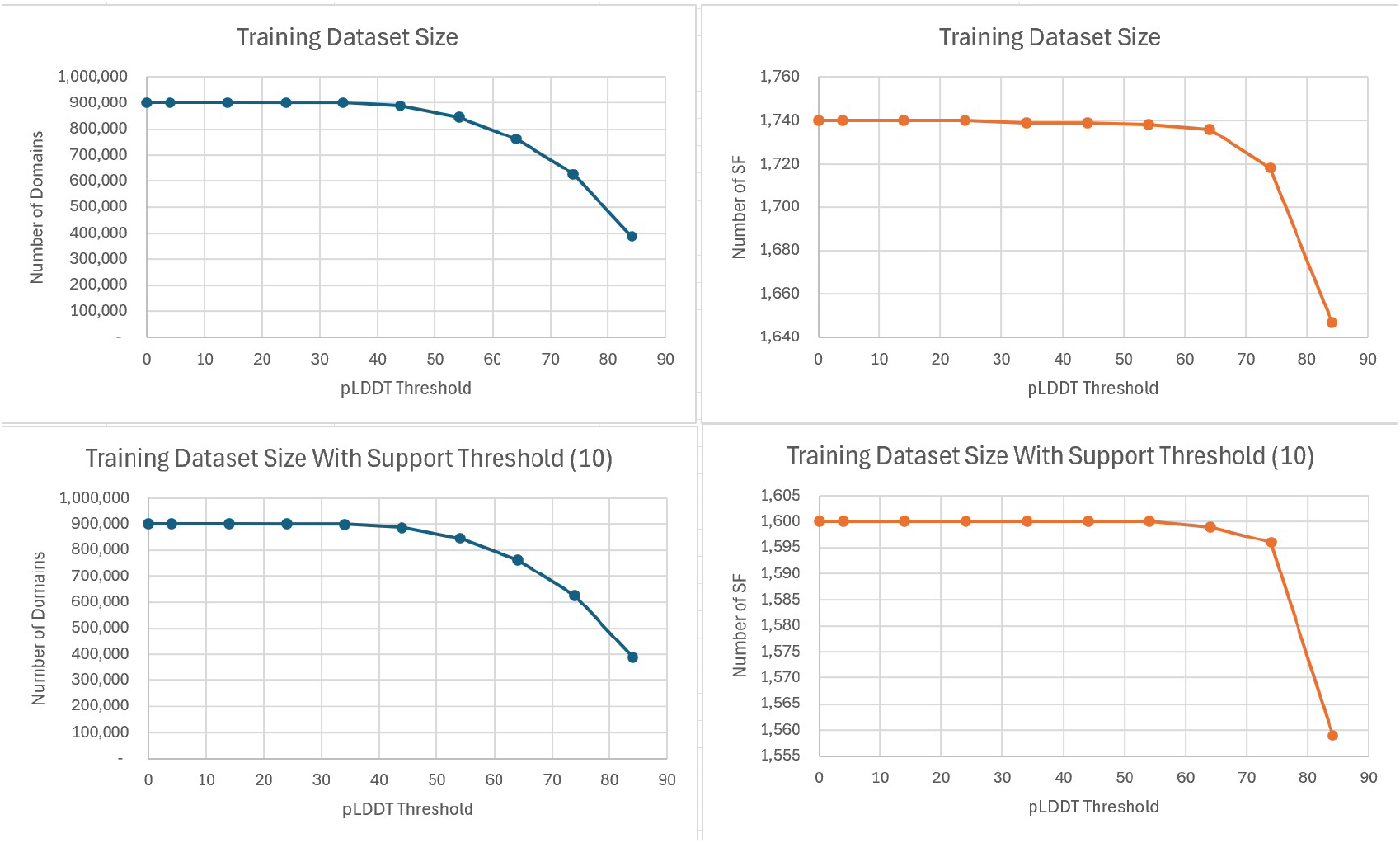
Training set size evolution with the different filters. These plots show training set size for the different combinations of thresholds for the pLDDT filter and the support filter tested for CATHe2. The size is either counted with domain number of SF number.

### 2.2. 3Di processing

Structural Alphabets (SA) are sets of clusters grouping similar 3D conformations of short strings of AA. To each cluster is associated a letter, the set of clusters constituting an alphabet. They are used to represent the full 3D shape of proteins in one dimension, by building a sequence of SA letters, allowing for various 1D processing of 3D information. Most SA, such as CLE [48], 3D-BLAST [49] and Protein Blocks (PB) [50], discretize the conformations of short stretches of usually 3–5 Cα atoms. The SA chosen for this study is called 3D interaction (3Di) [18], it does not describe the backbone but rather tertiary interactions. The 20 states (or 20 letters) of the 3Di alphabet describe the geometric conformation of each residue, with its spatially closest residue. 3Di has three key advantages over traditional backbone structural alphabets. First, a weaker dependency between consecutive letters, secondly, a more evenly distributed state frequencies, both enhancing information density and reducing false positives, thirdly, the highest information density is encoded in conserved protein cores and the lowest in non-conserved coil/loop regions, whereas the opposite is true for backbone structural alphabets. In CATHe2, 3Di are computed using Foldseek [18] on PDB files from the RCSB Protein Data Bank (PDB) [51] for the test set and validation set. As the training set is composed of domains from the AlphaFold Protein Structure Database (AFDB) [52], Computed Structure Models (CSM) PDB files are extracted from there to compute 3Di sequences for this dataset. See Figure 4 for the whole CATHe2 input creation process.

**Figure 4:**
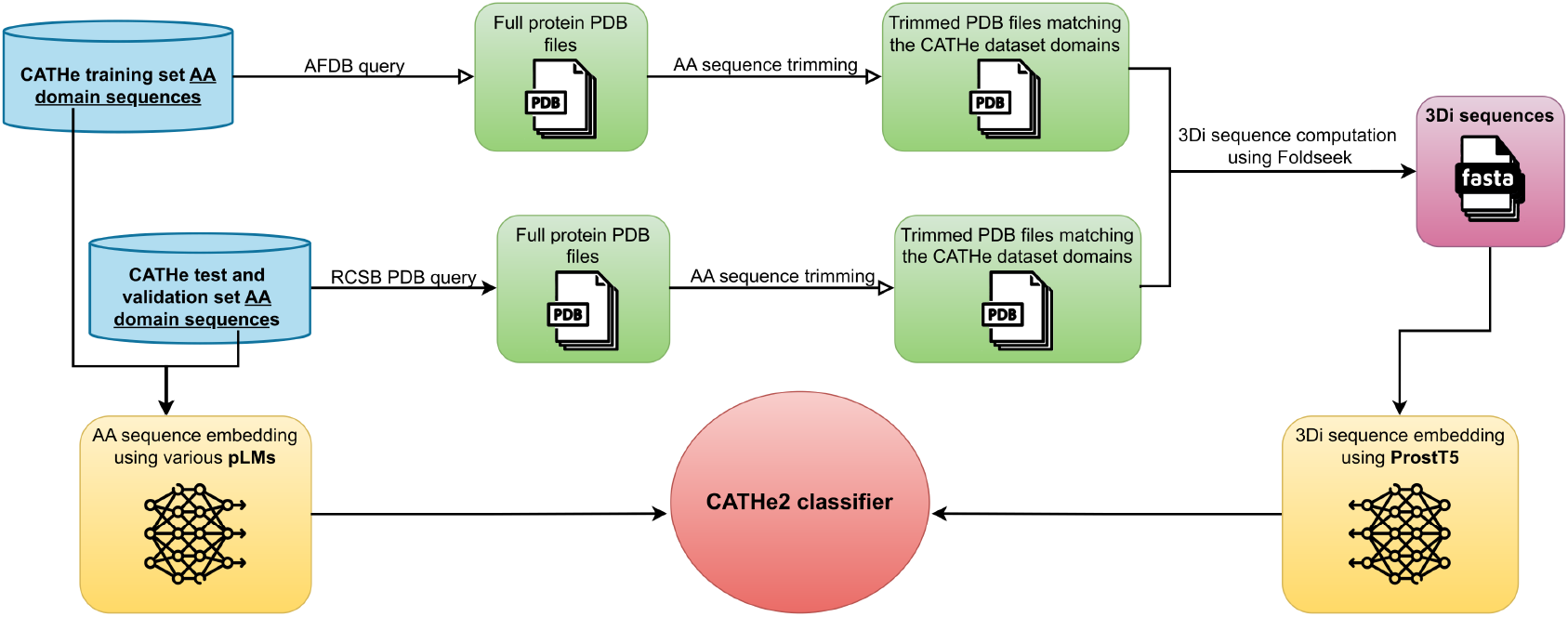
AA and 3Di sequence processing flowchart. CATHe datasets are composed of protein domains, not full protein sequences, therefore these domains had to be aligned to PDB and AFDB pdb file sequences, then trimmed, in order to keep only the 3D structure corresponding to the CATHe dataset domains. For this alignment, the Pairwise Aligner method from the Bio library was used [53]. These trimmed 3D structures were then used to compute 3Di sequences for ProstT5 to transform into 3Di sequence embeddings. Finally, combinations of 3Di embeddings and AA sequence embeddings were tested to train the CATHe2 classifier in order to get the highest F1 score possible. Meanwhile other hyperparameters than CATHe2 input type were tested like dataset filter values or classifier architecture.

### 2.3. Models

CATHe2 is an FNN classifier for which 7 hyperparameters were fine tuned to achieve the best results possible. First, the number of layer blocks; a layer block referring to a sequence of adjacent layers in this order: dense layer, LeakyReLu layer (Leaky Rectified Linear Unit layer), batch normalization layer, dropout layer. The other hyperparameters were the dropout rate for the dropout layer, the dense layer size, the pLDDT threshold, the support threshold, the input type (AA embeddings only, 3Di embeddings only or the concatenation of both) and finally, the pLM used to embed AA sequences, which are presented in this section. It is to note that apart from ProtT5, the pLMs described below are all recent and were launched after the publication of CATHe paper.

- ProtT5 (i.e, ProtT5-XL-U50) is the medium sized T5 encoder and the best-performing one from the ProtTrans paper [15]. This model is purely AA sequence based, meaning no structure information was used to train it. It was trained on BFD100 [47] and fine-tuned on Uniref50 [54]. Although ProtT5 embeddings were already used in CATHe, it is tested again for CATHe2 with a different code for generating embeddings, a better fine-tuned classifier architecture alongside 3Di embeddings in input. In this study ProtT5 proves again to be an excellent pLM for protein feature extraction by yielding very good results despite being rather old (2020).
- The ProstT5 pLM is a fine-tuned version of ProtT5 trained on both 3Di and AA span denoising task and bi-directional translation [27]. As a result, the encoder of ProstT5 can produce embeddings for both AA and 3Di sequences. The 3Di sequences used to train ProstT5 were computed from protein structures in AFDB [27, 55]. ProstT5 is already being used in many works, often for 3Di prediction from primary sequences, bypassing full 3D prediction [56, 57]. Moreover, in the ProstT5 paper, a CATH annotation benchmark is also carried out, using EAT and EAT+CL between embeddings from various inputs (AA, 3Di, ProstT5 predicted 3Di (p3Di), and the concatenation of AA and p3Di) comparing results with various pLM embeddings. The best results being consistently found to be with the ProstT5 pLM. In the CATHe2 code ProstT5 is called ProstT5 full as a half precision version called ProstT5 half is also tested for AA embedding computation (float16 weights instead of float32).
- TM-Vec is part of the tool duo TM-Vec+DeepBLAST whose purpose was to enable faster remote homology detection using structural alignment and deep learning [58]. TM-Vec produces structure-aware protein sequence embeddings, designed to help predict the TM-score [59], a measure of structural similarity between two protein sequences without the intermediate computation of their structures, even for remote homologs that fall below the 10% sequence identity. The model was trained on sequences from the CATH and SwissModel [60] structural databases. This model is actually based on ProtT5, with extra layers to help predict TM-scores [58, 55]. For their CATH annotation benchmark, the TM-Vec team used EAT to find that TM-Vec separates CATH structural classes more clearly than the default ProtT5. Their results show that across each level of the CATH hierarchy, TM-Vec outperformed FoldSeek [18], MMseqs2 [19] and finally ProtTucker [25] which combines ProtT5 embeddings, EAT, and Contrastive Learning, for CATH annotation inference with HBI [25].
- ESM2 was introduced in 2023 alongside the ESMFold tool that predicts protein structure from AA sequences only [45]. ESM2 is a purely AA sequence based pLM with an attention mechanism to learn pairwise interactions between amino acid sequences. It was trained on the Uniref50 and Uniref90 databases to learn large amounts of information and representations from protein sequences using masked language modelling (MLM). The same model was trained on multiple scales, ranging from 8 million parameters to 15 billion parameters. It is the 15 billion parameter version that was tested in this study (also called ESM2 15B) as it was shown to perform better overall [45, 55]. To fit this large model into the available memory space for CATHe dataset embedding, only a half precision version of ESM2 15B was tested (float16 weights instead of float32). This pLM was included in CATHe pLM tests as it was a recent pLM that had been shown to capture the primary sequence features related to 3D conformation. Plus it is very large and the proportionality between language model size and the richness of its learned representations has been validated in many fields [23], however, in this context, larger is not always better [15]. Even if ESM2 15B had shown incredible results, which is not the case (see Section 3), using it for the final inference model of CATHe2 would not have been practical due to the enormous amount of memory required just to load it, even in half precision. To use such heavy pLMs in regular settings, memory efficient techniques have to be employed like the ones used in ESME [61].
- Ankh is a purely AA sequence based pLM using a T5like architecture with a 48-layer transformer trained with MLM on UniRef50 sequences [23, 62, 11]. According to the corresponding paper, Ankh is a pLM that focuses on protein-specific optimization rather than relying on large model size, i.e, involving empirical, knowledge-guided optimization to improve performance without a proportional increase in computational resources. There are actually two Ankh models that were tested in this project, Ankh large, which is the larger version of Ankh, and Ankh base which is smaller. Ankh was also benchmarked on a CATH annotation task and yielded rather good results compared to ESM2 and ProtT5, outperforming them for the mean accuracy across all CATH levels [23].

## 3. Results

In order to have a comprehensive evaluation of different pLMs with different hyperparameters, we have run many preliminary experiments on CATHe’s large dataset in order to come up with the ideal combination. To do this, some comparisons have been made with fixed classifier structure and dataset filters to compare all pLMs, and ProstT5 full was selected to fine tune other hyperparameters on (fine tuning all hyperparameters for each pLM would have been too computationally costly). Then a grid search has been performed to approach the best classifier structure and dataset filter values, guided by training loss analysis. Some more insights for this fine tuning and for the preliminary experiments are available in the supplementary material to this paper.

Based on the above, we have chosen the best combination found of all hyperparameters and filters to present a summary of the results. Although balanced accuracy and MCC were also computed for CATHe2 models, we have chosen to focus on F1 score since the dataset is highly unbalanced (95% confidence windows were computed using bootstrapping on the CATHe test set with 1000-folds and the standard deviation from this bootstrapping method was then multiplied with 1.96 to obtain the 95% confidence interval values). All F1 scores for all models tested for CATHe2 are available in the /results/perf dataframe.csv file in the GitHub project. Table 1 presents those results on the CATHe large dataset.

**Table 1:** CATHe2 final models performance.

| Embedding input for CATHe2 | F1 score |
| --- | --- |
| ProstT5_full | **82.3 ± 1.3%** |
| ProstT5_half | 81.9 ± 1.3% |
| ESM2 15B | 80.6 ± 1.3% |
| ProtT5 | 80.6 ± 1.3% |
| Ankh_large | 78.3 ± 1.4% |
| Ankh_base | 77.3 ± 1.3% |
| TM-Vec | 76.5 ± 1.4% |

Hyperparameters corresponding to these results are the ones used for the best CATHe2 model i.e., the ProstT5\_full embedding based model on the large dataset without support threshold. These hyperparameters are Nb\_Layer\_Block: 2, Dropout: 0.3 Input\_Type: AA+3Di, Layer\_size: 2048, pLDDT\_threshold: 24.0.

These results suggest ProstT5 full >ProstT5 half >ESM2 15B >ProtT5 >Ankh large >Ankh base >TM-Vec in the CATHe2 context, but it is important to note that each model and version of models can perform differently relative to the others, based on the other hyperparameters. Moreover, comparing the performance of different pLM embedding inputs in the same ANN classifier like in Table 1 is questionable as each pLM produces embeddings which are of different sizes and probably need different ANN classifier architecture to show maximum performance. However, ProstT5 full AA+3Di embedding based version of CATHe outperforms significantly other CATHe2 models and previous models, i.e., CATHe (F1 Score of 82.3% Vs 72.4%) [14]. As shown in Table 1, the fourth row, even with ProtT5 using our improved ANN algorithm and adjusted hyperparameters, the F1 Score didn’t exceed 80.6%, which shows the added value of ProstT5 over ProtT5. The ProstT5 full embedding, AA only model available for inference in CATHe2 has the following hyperparameters: Nb Layer Block: 2, Dropout: 0.1 Input Type: AA only, Layer size: 1024, pLDDT threshold: 0, Support threshold: 0, and has an F1 score of 78.7% ± 1.3 on CATHe large dataset. The introduction of 3Di embeddings in addition to AA embeddings improved classifier performance whereas 3Di embeddings on their own clearly underperform. This implies that AA sequences have a lot more useful information for CATHe2 classifiers than 3Di sequences. This was expected since the classification of CATH’s superfamilies is mainly based on sequence similarity, that is, evolutionary information. Not to add that the 3Di SA, similar to other structural alphabets, does not fully capture the structural information, in other words, it is not a one-to-one relationship (one SA sequence could correspond to more than one structure).

In this section, we will present the results of the advanced and further experiments, i.e., the pLDDT and support filters. As shown in Figure 5 and as mentioned earlier, the combination of 3Di and AA inputs yielded consistently better results than AA only trained models for the same hyperparameter combination, especially when applying a pLDDT threshold around 24, whereas 3Di only models performed much worse consistently even with a high pLDDT threshold. Surprisingly, high pLDDT thresholds resulted in poor F1 scores. The reason behind this is the loss of many domains in the training dataset. When applying a ‘strict’ pLDDT filter (84 for example), the size of each class/SF decreases significantly, which naturally affects the accuracy of the model. The conclusion here is that the negative effect of decreasing the size of the training dataset outweighs the positive effect of removing domains whose 3D structure is of medium quality. This was also expected, because as shown in Figure 5 (AA+3Di Vs AA Vs 3Di), the values that the 3Di has added is relatively quite small, becauseagain – superfamily annotations are heavily based on evolutionary information, consequently, one wouldn’t expect a tangible improvement when removing ‘accurate’ AA sequences whose 3Di sequence is not ‘that accurate’. As for the second filter, i.e., the support filter, results are shown in Figure 6. due to a higher number of SF lost with support threshold (172 SF were lost with a threshold equal to 10), no models trained with it made it to the final version of CATHe2 despite achieving a very high performance of 88.0 ± 1% F1 score and accuracy of 93.8 ± 0.6% on the large dataset. This result shows again that training data sample size is very important to improve model performance and implies that a good part of future development should be focused on the gathering of larger well curated datasets.

**Figure 5:**
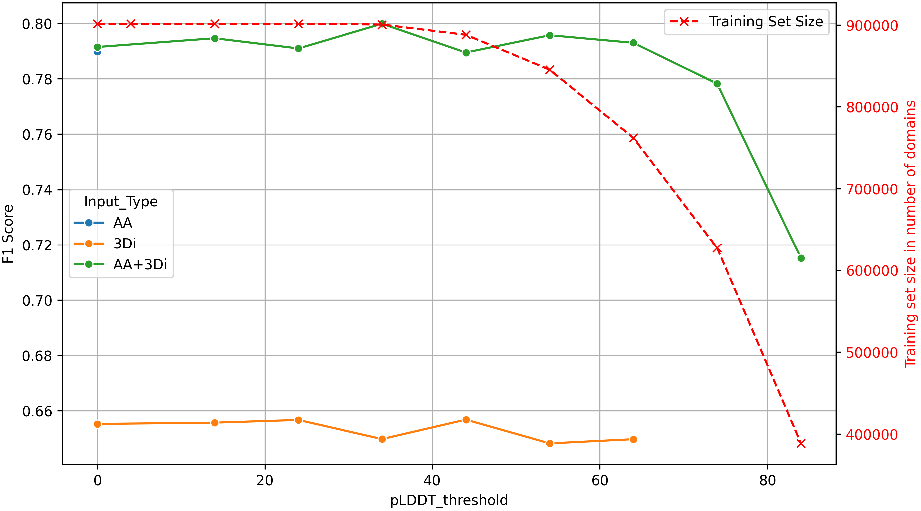
CATHe2 F1 score input type comparison along pLDDT threshold for a ProstT5 model. For AA only input there is only a single data point, for pLDDT threshold 0, because pLDDT is a 3D CSM confidence score, whereas no 3D models are used with only AA inputs. Moreover, the training set size mentioned for this AA only data point is not the one mentioned by the second y axis, but rather the full initial training dataset containing 1,039,135 domains. Hyperparameters for this model comparison are model embedding: ProstT5 full, Support threshold: 0, Nb layer block = 2, Layer size = 1024, Dropout = 0.3. Results are taken from CATHe2 large dataset, with no support threshold.

**Figure 6:**
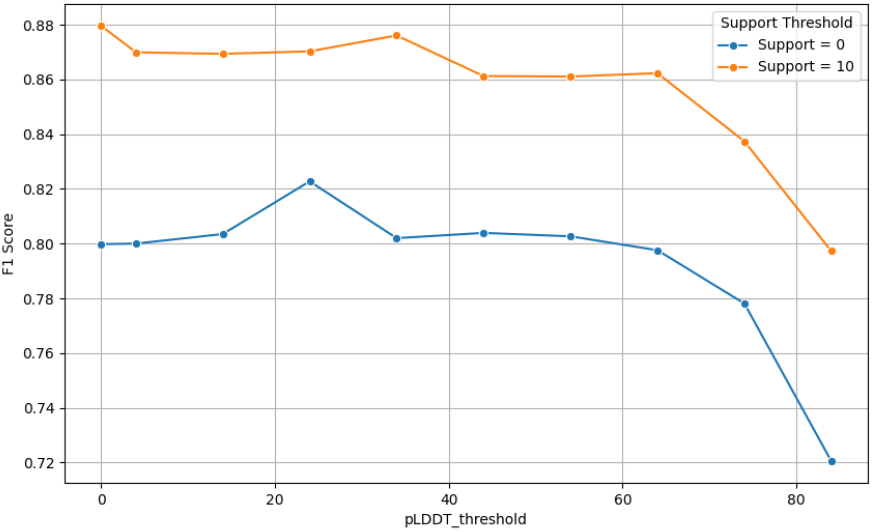
F1 score comparison for support threshold for the best hyperparameter combination. Contrary to the first plot, the hyperparameters here correspond to the best combination found for CATHe2 on its large dataset, i.e ProstT5 full AA+3Di embeddings with Dropout = 0.3, Layer size = 2048 and Nb layer block = 2

Compared to the previous ANN classifier architecture, CATHe2 best models have more layers (2 layer blocks instead of 1) and larger dense layers (2048 nodes instead of 128). This increase of classifier complexity has been shown to improve results up to a certain point.

To conclude on CATHe2 results, Table 2 presents a comprehensive performance review of pLM and HBI technique combinations for CATH annotation inference ranked from best to worst, according to the literature found on this topic. The performance metric here is the mean accuracy across all CATH levels. (When no input type is mentioned, it means only AA embeddings were used). This ranking is up to discussion as the accuracy is not so relevant for unbalanced datasets, F1 score would be more informative. Moreover (apart from CATHe and CATHe2 which predict the whole CATH annotation, i.e, the SF annotation) these are averages over multiple predicted CATH level annotation, plus each result relies on different datasets across multiple papers. Finally, the same technique with the same pLM trained and tested on the same datasets can yield varying results across replications with sometimes as much as a 7% standard deviation. Still, this ranking of HBI techniques for CATH annotation tends to show that classifiers >EBA >EAT+CL >EAT for CATH annotation transfer. This is also up to discussion as each technique has different variants depending on input type, distance metric, alignment algorithm, etc. Moreover, each technique requires a different computation time, whether it is for model construction or inference on new data. This general ranking is nonetheless supported by other papers [63]. From this paper review, classifiers seem to perform better at CATH HBI, and ProstT5 appears to be particularly effective for CATH annotation transfer. Note that the good placement of CATHe and CATHe2 classifiers could be explained by the fact that they are restricted on around 1700 CATH annotations only, whereas other HBI techniques can query a larger set of possible annotations.

**Table 2:**
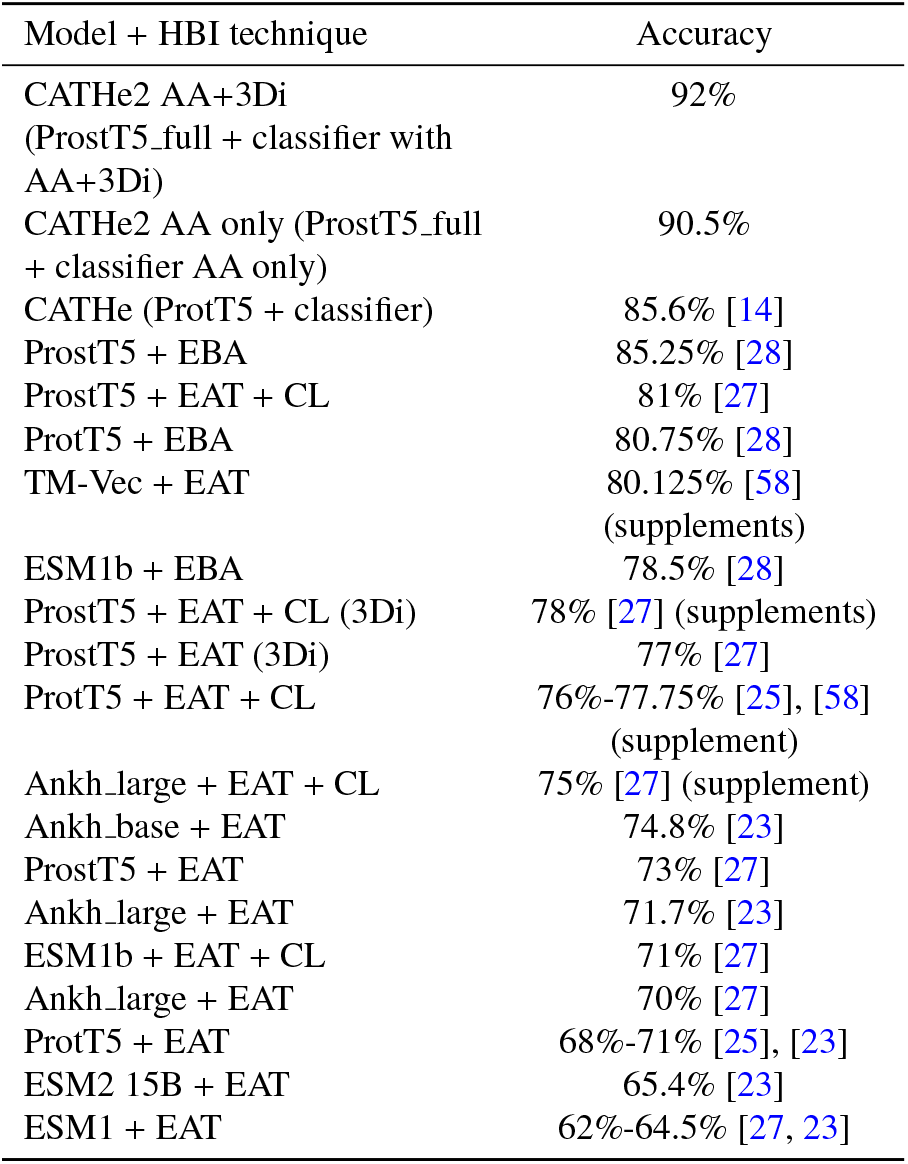
CATH annotation HBI technique performance ranking.

## 4. Discussion

### 4.1. Critical Analysis of the Study

In this paper, we proposed CATHe2, an updated version of CATHe (the CATH superfamilies automated classifier produced by the CATH team). Using roughly the same large dataset, we were able to attain an F1 score of 82.3% vs 72.4% (accuracy of 92.2% vs 85.6%). After a thorough study and comparison using different pLMs, hyperparameters, and methodologies, we propose the CATHe2 classifier with 2 layer blocks, 0.3 dropout rate for the dropout layers, and 2048 nodes for the dense layers. The input of this final model is a concatenation of ProstT5 AA sequence embeddings and ProstT5 3Di embeddings (pLDDT threshold of 24 for the CSM used to compute 3Di sequences). Although the CATHe2 dataset is quite large and is representative of the most populated SFs (The 200 most highly populated superfamilies in CATH alone represent 62% of AF2 domains [7]), CATHe2 is bounded to infer only CATH superfamilies present in these ~ 1700 SF, and therefore cannot predict a superfamily that was not in the training set, in particular, it lacks the SF labeled ‘other’ present in former CATHe. The absence of this class could explain the performance improvement of CATHe2 over CATHe, but it is likely to explain only a small part of this improvement as a ProtT5 model trained with the same hyperparameter as CATHe, without the ‘other’ SF has only an F1 score of 73.8 ± 1.3%, compared to the 72.4 ± 0.7% of CATHe and the 82.3 ± 1.3% of the final CATHe2 model. Beyond the removal of the ‘other’ superfamily detection, we had to remove 32 SF (out of 1773), due to the absence of their corresponding structures, missing those SF from the model is likely to explain only a small part of CATHe2 F1 score improvement too. Indeed, a classifier similar to the CATHe one, trained on AA sequences only, but without those 32 SF (without the ‘other’ class), improved by 1.5% only compared to the same model but trained with those SFs (still without the ‘other’ class).

### 4.2. Avenues for improvement

In this part, many new techniques and avenues for improvements are proposed to help future research on CATH annotation automation. A first promising improvement lead would be to implement Light Attention [63] when computing pLM embeddings per protein, instead of the average pooling on residue embeddings like in CATHe and CATHe2. As always, training the classifier with a larger, well curated dataset, is also expected to improve results, to make this easier, 3Di sequences could be inferred from AA sequence only using ProstT5 associated CNN [27] or ESM2 embeddings [41]. Training the pLMs again during classifier training is another promising lead. It may orient embedding computation toward the specialized task of CATH annotation prediction and thus improving results [64], although back propagation training for specific task fine tuning is not always successful and can lead to catastrophic forgetting or overfitting [15, 65]. Not to mention that such task is computationally very heavy for large models, especially without using complexity reduction methods like reduced precision or low ranking adaptations (LoRA) [66, 67, 68, 69]. For CATHe2 the concatenation of AA embeddings from one pLM and 3Di embeddings from ProstT5 was tested. To go further, multiple AA embeddings from multiple pLMs and the 3Di embeddings from ProstT5 could be concatenated (or even 3Di embeddings from multiple 3Di trained pLMs). Specially, concatenating AA embeddings from ProstT5 and ProtT5 as the ProstT5 paper suggests, could improve performance [27]. Although 3Di sequences were used in this study, it may be interesting to test protein blocks (PB) [50] instead as the 3Di alphabet was not developed to maximize remote homology search sensitivity, but to model tertiary interactions in protein structures [18]. Even when using 3Di, a bilingual model like ProstT5 may not be optimal. Thus, a potential improvement on CATHe2 would be to use AA sequence embeddings from an AA sequence specialized pLM like ProtT5 and a 3Di (or PB) sequence specialized pLM which, to our knowledge, has not been developed yet. The pLMs tested for CATHe2 were just a promising subset of all current pLMs suited for the task. Other promising pLMs exist, ProSST for instance is a good candidate, as it is a hybrid AAstructure pLM that produces its own 3Di like structural representation with an advanced structure quantization method and a better attention formulation to leverage the structure cues [70]. Finally, SaProt embeddings could be used to improve CATHe2 results too. Very similar to ProstT5, SaProt is based on ESM2 650M instead of ProtT5. For SaProt, AA and 3Di were fused to create a novel structural alphabet on which it was trained [71]. SaProt could be one of the best pLM to introduce structural information as suggested by other researchers [38]. Many other pLMs could be tested, extensive lists can be found in the literature [11, 12]. Lastly, other avenues for improvement include using embeddings from another hidden layer than the last one, using embedding denoising [10], and using other classifier architectures like CNN classifiers or even a combination of CNN classifiers, which seem to be more efficient than the simple FNN classifier used for CATHe and CATHe2 [72].

## Acknowledgments

Orfeu Mouret would like to thank Jad Abbass and Kingston University staff in general for their support.

## CRediT authorship contribution statement

Orfeu Mouret: Methodology, Software, Data curation, Visualization, Investigation, Writing-Original draft, Conceptualization. Jad Abbass: Conceptualization, Supervision, Writing-Reviewing and Editing.

## Declaration of Generative AI and AI-assisted technologies in the writing process

During the preparation of this work the author(s) used ChatGPT-4o in order to assist with phrasing and to correct spelling. After using this tool/service, the author(s) reviewed and edited the content as needed and take(s) full responsibility for the content of the publication.

