## Supplementary_material for "CATHe2: Enhanced CATH Superfamily Detection Using ProstT5 and Structural Alphabets"

**Outline:**
This supplementary document provides more details on CATHe2 preliminary experiments, hyperparameter fine tuning and training set modifications.

### Preliminary Experiments

The first step of CATHe2 grid search fine tuning was to identify the "best" pLM to embed AA sequences with. This "best" pLM depends on the hyperparameters of course, that's why a rather large range of hyperparameter combinations have been tested to try selecting the pLM with the most potential for further fine tuning. The next plots show some performance comparisons made during this step of preliminary experiments (during this first step the best combination of hyperparameters for ProstT5 full was obviously not found yet). The next step of preliminary experiments was to begin revealing clues about what the best values were for each hyperparameter, the next plots were also used to this effect.

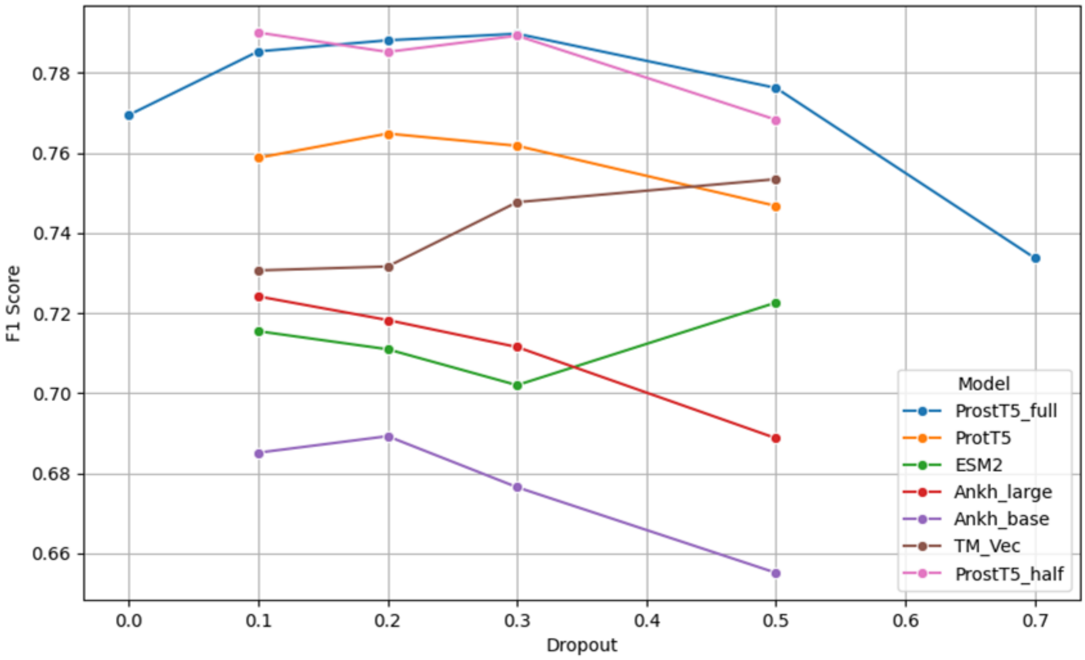


Hyperparameters: large dataset (is_top_50 = False)
pLDDT threshold: 0
support threshold: 0
input type: AA
Nb layer block: 2
Layer size: 1024

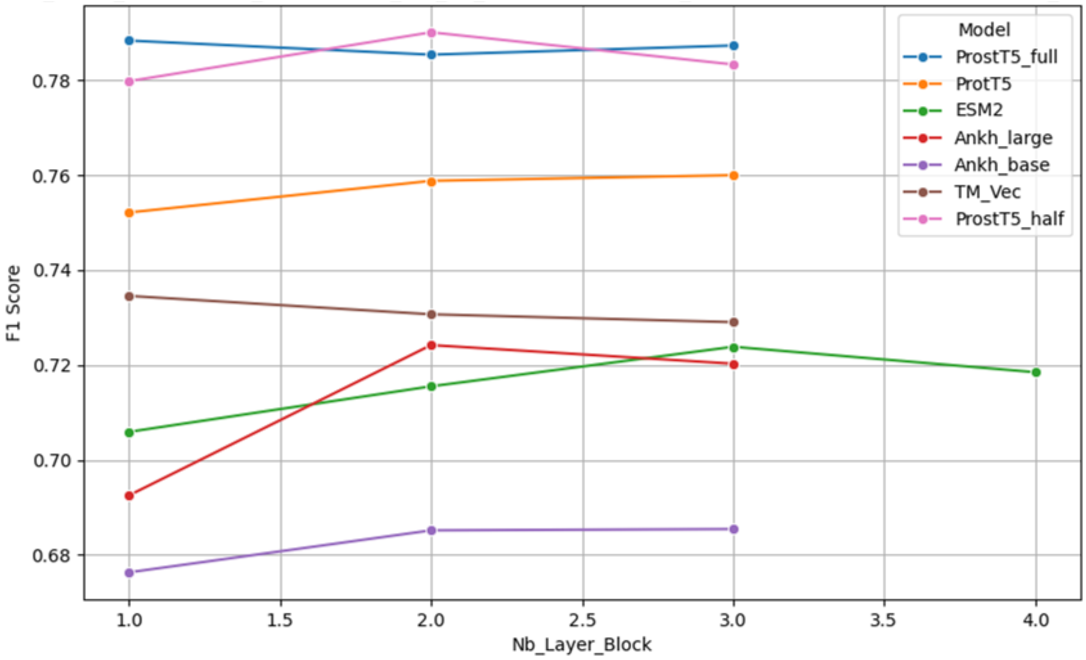


Hyperparameters: large dataset (is_top_50 = False)
pLDDT threshold: 0
support threshold: 0
input type: AA
Dropout: 0.1
Layer size: 1024

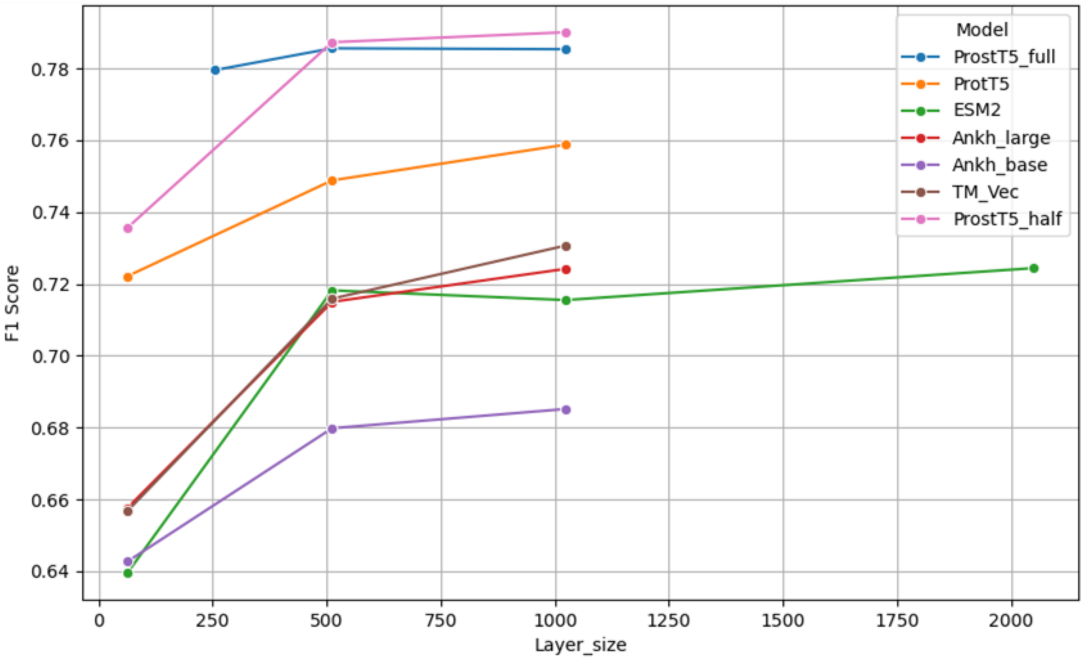


Hyperparameters: large dataset (is_top_50 = False)
pLDDT threshold: 0
support threshold: 0
input type: AA
Dropout: 0.1
Nb layer blocks: 2

### Hyperparameters fine tuning

**Loss curve analysis**

As mentioned in the paper, the fine tuning of CATHe2 was conducted using a grid search guided by loss curve analysis. During CATHe2 model training, training loss and validation loss were saved to guide the next step of grid search. Here is an example of such a curves on a plot (this one is the loss curve of the final ProstT5 full model).

Much

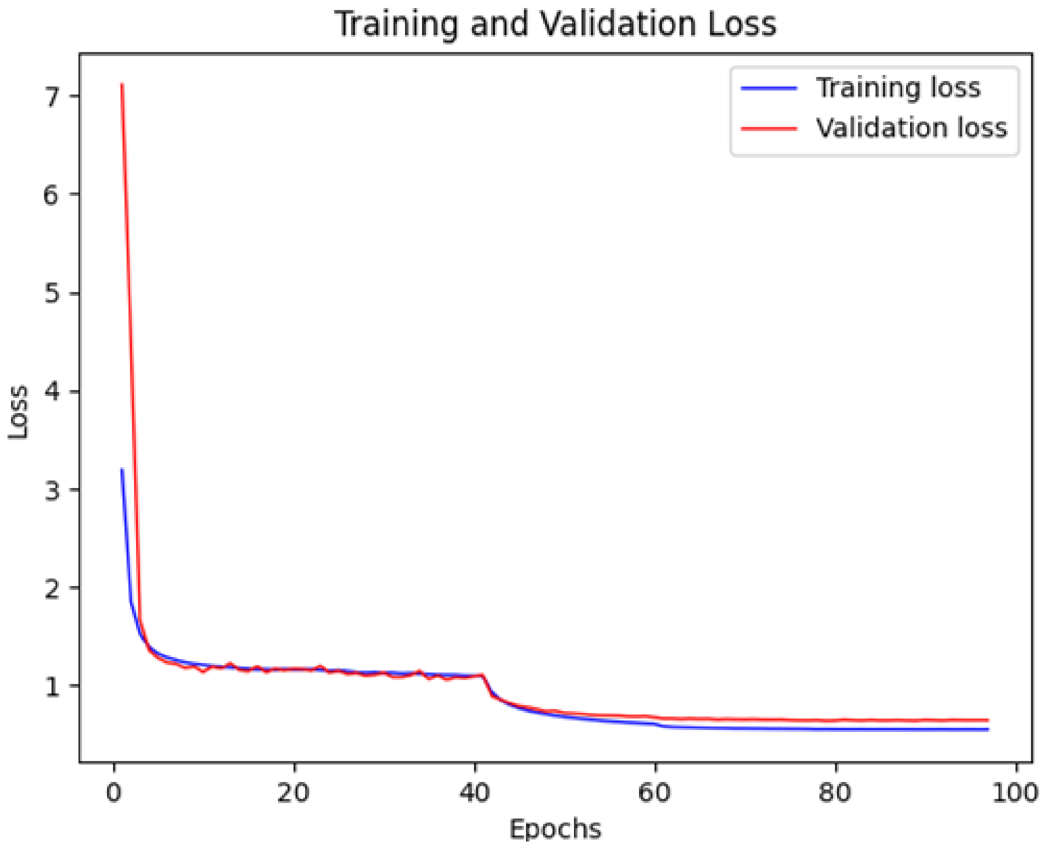


information can be deduced on such a plot. For instance an erratic loss curve can mean a too high learning rate. A validation loss getting close to training loss then separating from it can mean overfitting, which can be reduced by shortening epoch number or increasing dropout rate. A validation loss never approaching training loss can mean that the model does not have enough epochs to learn, or that the training set does not contain enough information to generalise, or that model structure is not complex enough to learn general classification rules. A too high training loss means that the model is not even learning classification rules for the training set, meaning this training set is not big enough or representative enough of classification rules, or that the model is not complex enough for example. All these clues and more can be derived from loss curves which gave some directions to explore more efficiently the hyperparameter space during grid search.

**Individual hyperparameter value analysis**

Model performance is linked to every single model hyperparameter in an unknown and probably non linear way. Thus trying to fine tune hyperparameters one by one is pointless, as for the same value of Dropout rate for example a large range of performance is possible depending on the other hyperparameters. All hyperparameters should be modified together to learn the right combination. However, it does not mean that nothing can be learned by analysing performance evolution when fixing all hyperparameters but one. It is expected for example that dropout rate should be increased when model complexity (number of layer blocks and dense layer size) is increased, to avoid overfitting. So learning the “right” dropout rate for a fixed model complexity can yield some indications about what is the “right” dropout rate to expect for other hyperparameter combinations. In addition, analysing performance along different hyperparameters, with all the other fixed can also yield clues about what performance to expect for other hyperparameter combinations, and what hyperparameter values are best. In the next plots for example, input type AA+3Di clearly performed better than any other input type. The following plots present some of the fine tuning process of ProstT5 full by showing performance evolution along a unique hyperparameter, while the others are fixed.

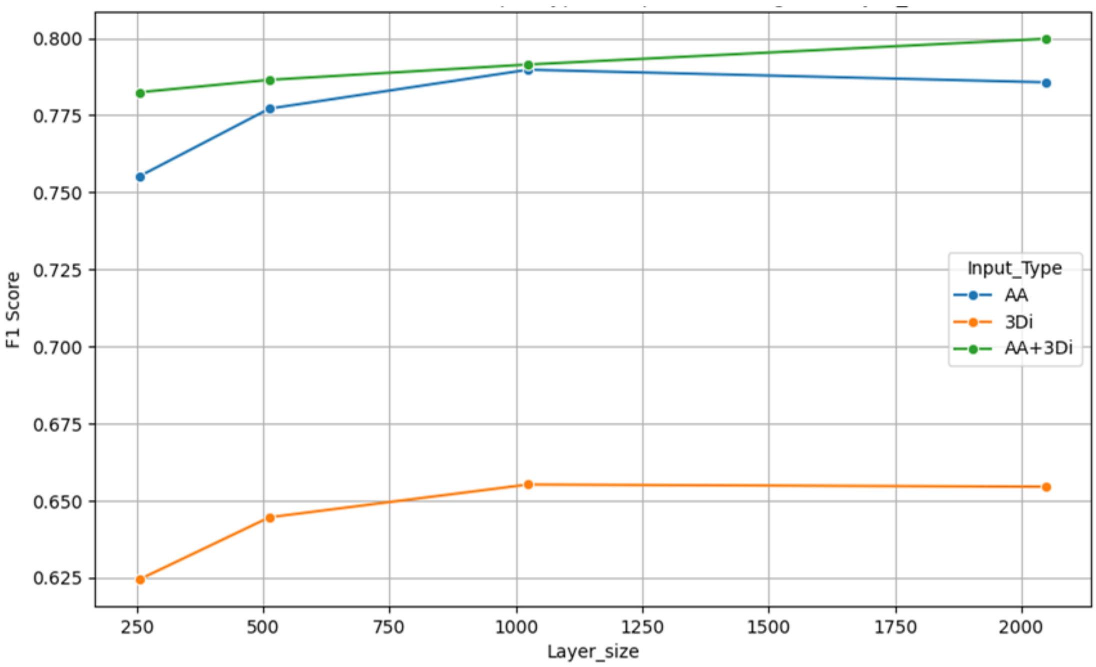


Hyperparameters: large dataset (is_top_50 = False)
pLM: ProstT5_full
pLDDT threshold: 0
support threshold: 0
Dropout: 0.3
Nb layer blocks: 2

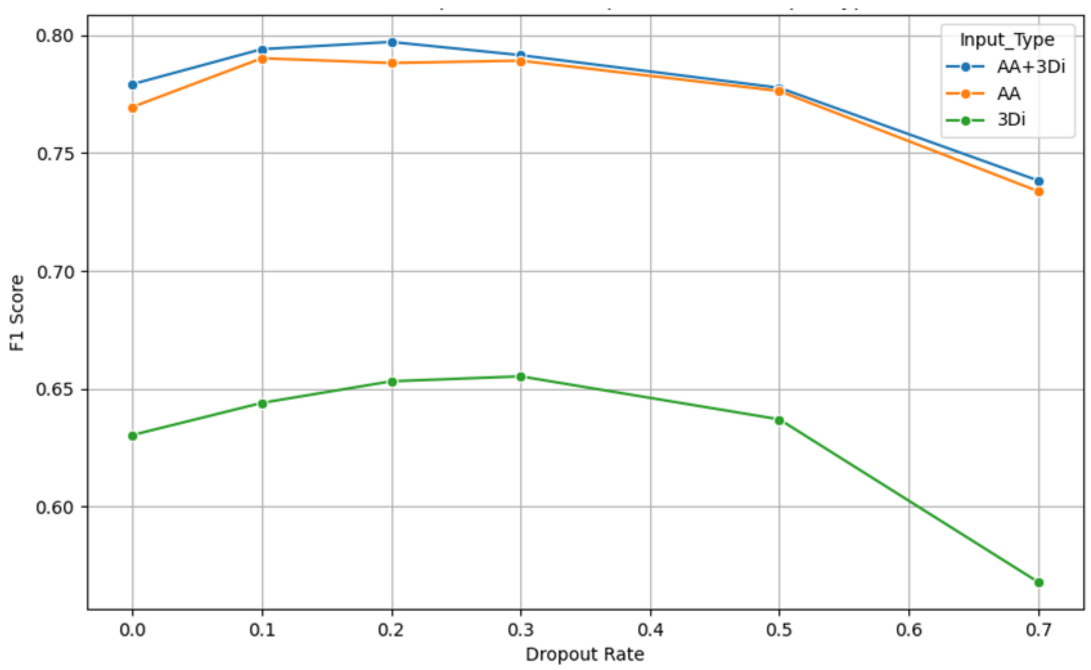


Hyperparameters: large dataset (is_top_50 = False)
pLM: ProstT5_full
pLDDT threshold: 0
support threshold: 0
Layer size: 1024
Nb layer blocks: 2

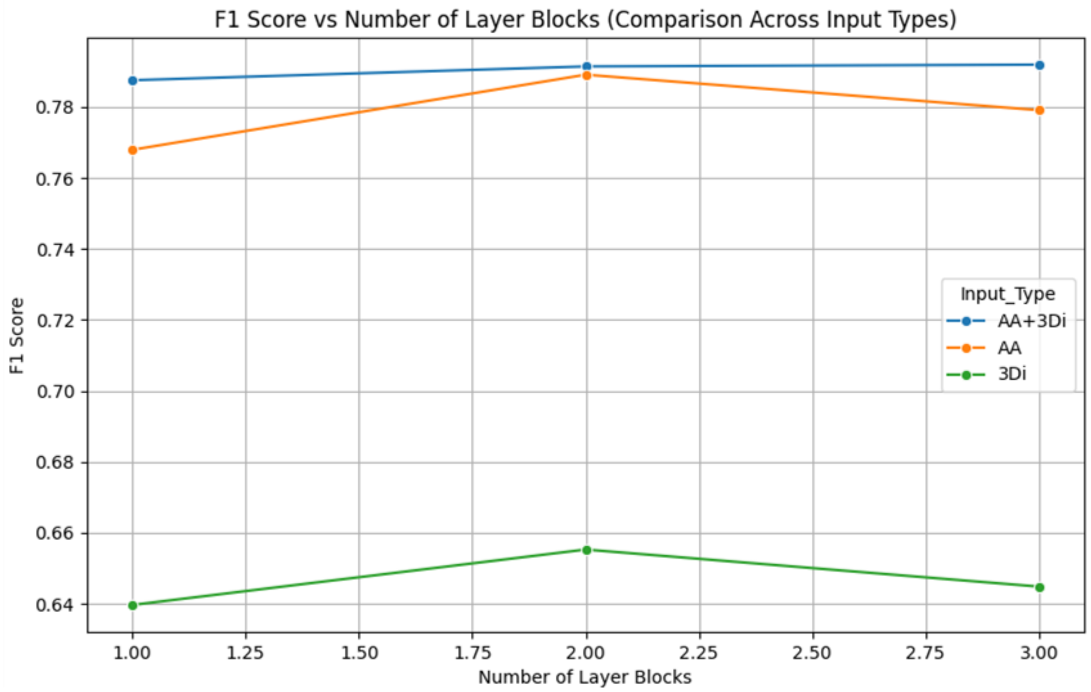


Hyperparameters: large dataset (is_top_50 = False)
pLM: ProstT5_full
pLDDT threshold: 0
support threshold: 0
Dropout: 0.3
Layer size: 1024

**Traininig set modifications relatively to CATHe2 filter thresholds**

| PLDDT threshold | Lost SF count | Number of SF remaining | Training set size | Lost domains Count | Top 50 filtering | Support threshold |
|---|---|---|---|---|---|---|
| 0 | 32 | 1,740 | 901,437 | 137,698 | FALSE | 0 |
| 4 | 32 | 1,740 | 901,437 | 137,698 | FALSE | 0 |
| 14 | 32 | 1,740 | 901,437 | 137,698 | FALSE | 0 |
| 24 | 32 | 1,740 | 901,429 | 137,706 | FALSE | 0 |
| 34 | 33 | 1,739 | 900,452 | 138,683 | FALSE | 0 |
| 44 | 33 | 1,739 | 887,929 | 151,206 | FALSE | 0 |
| 54 | 34 | 1,738 | 845,285 | 193 850 | FALSE | 0 |
| 64 | 36 | 1,736 | 762,164 | 276,971 | FALSE | 0 |
| 74 | 54 | 1,718 | 627,412 | 411,723 | FALSE | 0 |
| 84 | 125 | 1,647 | 388,478 | 650,657 | FALSE | 0 |
| 0 | 1,722 | 50 | 456,585 | 582,550 | TRUE | 0 |
| 4 | 1,722 | 50 | 456,585 | 582,550 | TRUE | 0 |
| 14 | 1,722 | 50 | 456,585 | 582,550 | TRUE | 0 |
| 24 | 1,722 | 50 | 456,585 | 582,550 | TRUE | 0 |
| 34 | 1,722 | 50 | 456,184 | 582,951 | TRUE | 0 |
| 44 | 1,722 | 50 | 449,897 | 589,238 | TRUE | 0 |
| 54 | 1,722 | 50 | 427,373 | 611,762 | TRUE | 0 |
| 64 | 1,722 | 50 | 384,638 | 654,497 | TRUE | 0 |
| 74 | 1,722 | 50 | 316,649 | 722,486 | TRUE | 0 |
| 84 | 1,722 | 50 | 192,677 | 846,458 | TRUE | 0 |
| 0 | 172 | 1,600 | 900,813 | 138,322 | FALSE | 10 |
| 4 | 172 | 1,600 | 900,813 | 138,322 | FALSE | 10 |
| 14 | 172 | 1,600 | 900,813 | 138,322 | FALSE | 10 |
| 24 | 172 | 1,600 | 900,805 | 138,330 | FALSE | 10 |
| 34 | 172 | 1,600 | 899,832 | 139,303 | FALSE | 10 |
| 44 | 172 | 1,600 | 887,321 | 151,814 | FALSE | 10 |
| 54 | 172 | 1,600 | 844,702 | 194,433 | FALSE | 10 |
| 64 | 173 | 1,599 | 761,630 | 277,505 | FALSE | 10 |
| 74 | 176 | 1,596 | 626,974 | 412,161 | FALSE | 10 |
| 84 | 213 | 1,559 | 388,204 | 650,931 | FALSE | 10 |
